# Potential limitations of micro-dystrophin gene therapy for Duchenne muscular dystrophy

**DOI:** 10.1101/2022.10.02.510519

**Authors:** Cora C. Hart, Young il Lee, Jun Xie, Guangping Gao, David W. Hammers, H. Lee Sweeney

**Author notes:** Corresponding author* **Correspondence:** H. Lee Sweeney, 1200 Newell Dr. ARB R5-216, Gainesville, FL 32610-0267. Equal contributions by these authors (Cora C. Hart, Young il Lee). Equal contributions by these authors (David W. Hammers, H. Lee Sweeney).

## Abstract

Adeno-associated viruses (AAVs) expressing versions of truncated dystrophin (micro-dystrophins) are being delivered at high doses to patients with Duchenne muscular dystrophy (DMD) in clinical trials. We examined this strategy with two different micro-dystrophins, similar to those currently in clinical trials, in a severe mouse model of DMD, the D2.*mdx* mouse, using doses of AAV comparable to those used in the clinical trials. We achieved high levels of micro-dystrophin expression in striated muscle with cardiac expression ∼10 fold higher than that observed in skeletal muscle. Significant, albeit incomplete, correction of the skeletal muscle disease is observed. Surprisingly, a lethal acceleration of cardiac disease progression occurs with one of the micro-dystrophins, while the second appears to benefit the heart. The detrimental impact on the heart in the first case appears to be caused by the high levels of micro-dystrophin in the heart resulting in competition between micro-dystrophin and utrophin at the cardiomyocyte membrane. While the significance of these observations for patients currently being treated with AAV-micro-dystrophin therapies is unclear since the levels of expression being achieved in the DMD hearts are unknown, it suggests that micro-dystrophin treatments may need to be carefully titrated to avoid high levels of expression in the heart.

## INTRODUCTION

Duchenne muscular dystrophy (DMD) is an X-linked disorder that affects approximately 1 in 5000 newborn males (1). It is the most common of the childhood muscular dystrophies and results from the lack of the membrane-associated protein, dystrophin, which is critical for proper force transmission in muscle cells (2, 3). The loss of dystrophin results in hypersensitivity to injury in the skeletal muscle and leads to cardiac dysfunction. The skeletal muscle initially undergoes rounds of injury and repair, but repair eventually begins to fail, and the muscles are replaced with fibrosis and fat. The muscle loss progresses from proximal to distal, with the loss of respiratory muscles and/or heart failure as the cause of death, generally in the second or third decade of life (4). The cardiac disease manifests first with diastolic dysfunction and later progresses to a dilated cardiomyopathy (DCM) and failure (5-8).

Gene therapy for DMD has entered the clinic in the form of several versions of a highly truncated dystrophin (micro-dystrophin) delivered via adeno-associated virus (AAV). While AAV is highly efficient at infecting and transducing striated muscle, its small packaging size (∼5 kb) makes it impossible to accommodate the full-length dystrophin coding sequence (∼14kb). This has necessitated using AAV to deliver the coding sequence of a highly truncated dystrophin (9, 10) or to use AAV to alter splicing of an out-of-frame dystrophin mRNA to create a deletion that restores the proper reading frame (11, 12). In either case, the goal is to express a truncated version of dystrophin in order to slow disease progression. This strategy essentially aims to transform DMD into a slower progressing muscular dystrophy, potentially more like some forms of Becker muscular dystrophy (BMD), which results from dystrophin mutations that create in-frame transcripts resulting in production of a variety of truncated forms of dystrophin that are associated with different rates of disease progression.

A number of questions surround the outcome of these trials, particularly the dosing and the potential efficacy of each of the different micro-dystrophin constructs currently in trial. It is unclear when and if there will be a need to redeliver the therapy if the skeletal muscle slowly turns over due to residual disease, resulting in eventual loss of the AAV DNA encoding the micro-dystrophin transgene. Thus, a major goal of the current micro-dystrophin trials is to express the micro-dystrophin at high levels throughout the skeletal and cardiac muscles, which potentially will limit the frequency of needing to redeliver AAV to the skeletal muscle. Since the cardiomyocytes do not turnover, redelivery will be unnecessary unless they were not adequately transduced with the first dose of virus.

Common features of the micro-dystrophin constructs currently in clinical trials (shown in **Figure 1**) include the N-terminal actin-binding region, four to five of the twenty-four spectrin-like triple helical bundles that make up the rod region, and a truncated C-terminus containing the β-dystroglycan binding site. Most of the preclinical work supporting the human trials has been performed using dystrophic mice of C57-based genetic backgrounds, which exhibit mild disease progression when compared to that of other mouse genetic backgrounds (13) and larger mammals (14, 15). While information concerning transgene delivery and expression can be gathered using these C57-based models, it is difficult to assess the true efficacy of AAV-micro-dystrophin gene therapies at correcting a severe, life-limiting striated muscle disease. Indeed, the lack of an animal model that is completely representative of the human disease has contributed to the discrepancy in results between preclinical and clinical research, and ultimately resulted in the termination of several DMD clinical trials (16). Therefore, in this study, we used a severe mouse model of DMD, the D2.*mdx* mouse harboring the *mdx* mutation on the DBA/2J genetic background (13, 17, 18) to evaluate the long-term impact of high dose AAV-micro-dystrophin on the heart and skeletal muscles in the face of a more aggressive disease progression.

**Figure 1.**
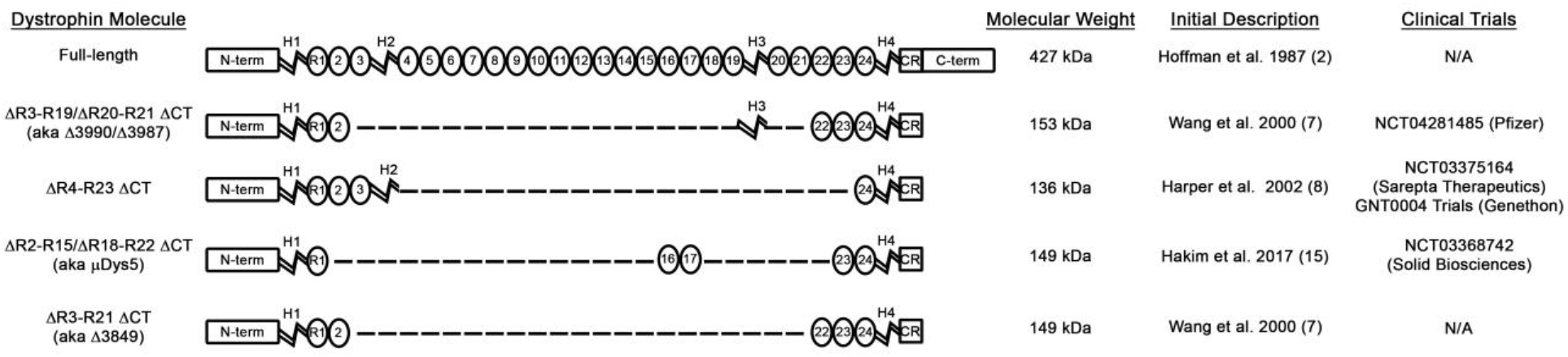
Structure of dystrophin and micro-dystrophin constructs. A schematic diagram of full-length dystrophin, the micro-dystrophin versions currently utilized in clinical trials. Also diagramed is the variant of the ΔR3-R19/ΔR20-R21 ΔCT micro-dystrophin construct used in this study (ΔR3-R21 ΔCT), which was compared to ΔR2-R15 ΔR18-R22 ΔCT.

We sought to compare two micro-dystrophin constructs that had not previously been compared, ΔR3-R19 /ΔR20-R21 ΔCT and ΔR2-R15 ΔR18-R22 ΔCT (see **Figure 1**). However, we could not achieve efficient packaging of the large ΔR3-R19/ΔR20-R21 ΔCT construct (3987bp) and instead used a smaller variant that differed only by the deletion of H3 (ΔR3-R21 ΔCT) to compare to ΔR2-R15 ΔR18-R22 ΔCT. In the initial publication describing the ΔR3-R19/ΔR20-R21 ΔCT and ΔR3-R21 ΔCT constructs, no significant difference was shown in efficacy (9). A more recent publication using a transgenic C57BL6.mdx mouse expressing the ΔR3-R21ΔCT construct showed rescue of skeletal muscle (19); therefore, we believe this to be an appropriate substitution for the evaluation of the long-term efficacy of clinically relevant micro-dystrophins. Using AAV doses similar to those used in current clinical trials, we find widespread transduction and sustained expression of both micro-dystrophins in skeletal and cardiac muscles of D2.*mdx* mice. Both therapies greatly slowed skeletal muscle disease progression, although not completely stopping it. The heart expressed higher levels of micro-dystrophin than did the skeletal muscle (∼5-10-fold higher). This cardiac overexpression accelerated the onset of a DCM, heart failure, and death in the treated mice with one of the micro-dystrophins tested (ΔR3-R21 ΔCT). The second micro-dystrophin (ΔR2-R15 ΔR18-R22 ΔCT) had little or no impact on the early disease progression in the heart, but prolonged median survival of the dystrophic mice to >18 months, at which time the mice developed DCM and progressed toward failure.

## RESULTS

### Clinical AAV doses enable widespread expression in D2.mdx striated muscle

We sought to evaluate how efficacious two different micro-dystrophins were at correcting the skeletal and cardiac muscle pathologies associated with the D2.*mdx* mouse model of DMD. In this long-term study, we utilized five-repeat microdystrophin constructs ΔR3-R21ΔCT and ΔR2-R15 ΔR18-R22 ΔCT. The ΔR3-R21ΔCT micro-dystrophin construct (also known as Δ3849 (9)) was chosen for this study because this version packages efficiently in AAV and has shown robust efficacy at correcting skeletal muscle when expressed as a transgenic (19). In a short-term assessment of ΔR3-R21ΔCT and ΔR3-R19/ΔR20-R21ΔCT we found that at equivalent expression levels, ΔR3-R21ΔCT better protected the skeletal muscle from the damage incurred with eccentric contractions (**Supplemental Figure 1**). The second micro-dystrophin construct evaluated, ΔR2-R15 ΔR18-R22 ΔCT, contains all of the elements of the construct in clinical trial.

As depicted in **Figure 2a**, this evaluation consisted of male D2.*mdx* mice receiving an intravenously (IV) delivered dose of AAV-packaged micro-dystrophin at 1 month of age. Both constructs were placed behind the CK8 striated muscle promoter (20) and packaged in AAVrh10 serotype vector, which has a high tropism for striated muscle (21) and is nearly identical to one of the vectors (AAVrh74) in clinical trials (22). AAV was administered systemically through the tail vein at a dose of 2×10^14^ gc/kg, which is similar to that currently used in the clinic (23).

**Figure 2.**
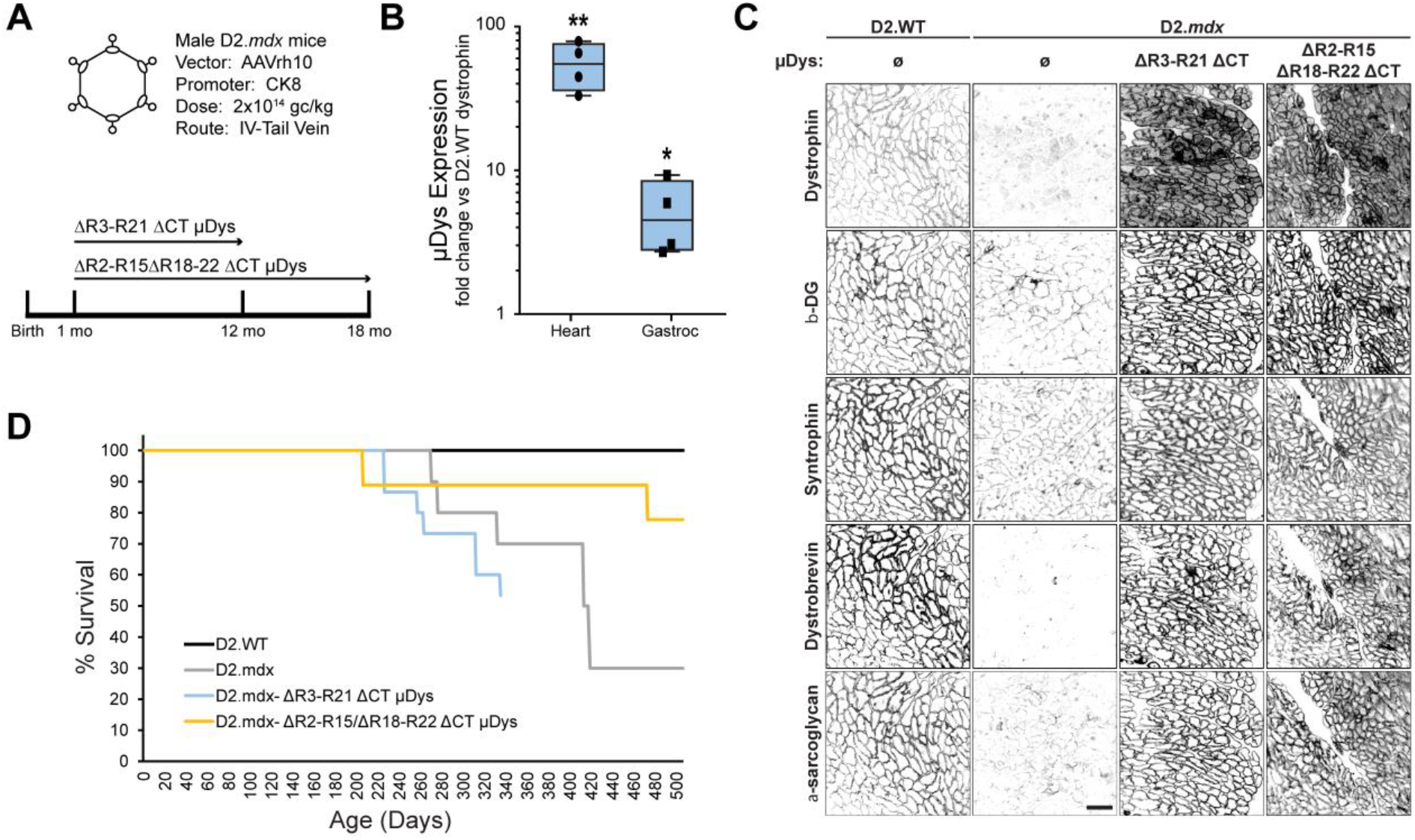
Adeno-associated virus-mediated expression of micro-dystrophin in striated muscles. (**a**) Male D2.*mdx* mice were injected with 2×10^14^ gc/kg of AAVrh10-CK8-micro-dystrophin (µDys) ΔR3R21ΔCT or ΔR2-R15 ΔR18-R22 ΔCT via tail vein at one month of age. (**b**) Administration of the 2 × 10^14^ gc/kg dose at 1 month of age leads to overexpression of µDystrophin compared to the levels of endogenous dystrophin in D2.WT mice: ∼55-fold and ∼5-fold over-expression in heart and gastrocnemius, respectively (n = 4; **p<0.01, *p<0.05; unpaired t-test vs endogenous dystrophin levels in gastrocnemius or heart). (**c**) Antibody-mediated labeling of heart transverse sections from D2.WT, D2.*mdx* and micro-dystrophin (µDystrophin)-treated D2.mdx mice reveals sarcolemmal localization of µDystrophin protein that mirrors sarcolemmal localization of full-length endogenous dystrophin protein in D2.WT cardiomyocytes. Scale bars: 100 µm. (**d**) Survival curve D2.WT, D2.*mdx* and micro-dystrophin treated D2.*mdx* mice reveals premature death of mice that received ΔR3R21ΔCT and an extension of life in mice that received ΔR2-R15 ΔR18-R22 ΔCT.

The treatment of D2.*mdx* mice in this manner resulted in equivalent striated muscle expression of both micro-dystrophins (**Supplemental Figure 2**). We observed robust and uniform expression of micro-dystrophin at the sarcolemma of cardiomyocytes, as detected by immunofluorescence (**Figure 2C**). The expression of micro-dystrophin in these tissues coincided with an increase in membrane-associated content of the dystrophin-glycoprotein complex (DGC) members β-dystroglycan, syntrophin, dystrobrevin, and α-sarcoglycan (**Figure 2C**).

Immunoblotting data estimate that the micro-dystrophin levels achieved by this treatment greatly exceed wild-type levels of native dystrophin in both the gastrocnemius and heart (**Figure 2B**; ∼5- and ∼55-fold greater, respectively). These results demonstrate that the treatment of D2.*mdx* mice with clinical doses of AAV-packaged micro-dystrophin leads to efficient transduction and micro-dystrophin expression in both skeletal and cardiac muscle. Despite equivalent micro-dystrophin levels being achieved by both constructs, we observed a striking difference in survival age between the two cohorts (**Figure 2D**). Whereas the ΔR2-R15 ΔR18-R22 ΔCT micro-dystrophin extends the average lifespan of D2.mdx mice, the ΔR3-R21ΔCT micro-dystrophin leads to a premature death. Therefore, the terminal measures for the mice treated with ΔR3-R21ΔCT or ΔR2-R15 ΔR18-R22 ΔCT were conducted at 12 and 18 months of age, respectively, as indicated **in Figure 2A**.

### Micro-dystrophin gene therapy partially corrects the D2.mdx skeletal muscle pathology

At terminal timepoints, *ex vivo* functional evaluations of diaphragm and extensor digitorum longus (EDL) muscles were performed. As anticipated by previous reports (10, 24, 25), micro-dystrophin treatment improved several features of skeletal muscle function including increases in diaphragm specific tension, EDL maximum force production, and EDL resistance to eccentric contraction-induced functional deficits, compared to untreated D2.*mdx* mice (**Figure 3A-D**). However, these functional improvements were, for the most part, significantly diminished compared to D2.WT values, with the diaphragm only achieving ∼50% of a full rescue effect (**Figure 3A**). In agreement with a partial skeletal muscle rescue by micro-dystrophin, the diaphragms of treated mice exhibited progressive fibrosis, albeit on a slower trajectory than those of untreated D2.*mdx* mice (**Figure 3G-H**). Therefore, systemic micro-dystrophin gene therapy does provide partial rescue of D2.*mdx* skeletal muscle. However, the resulting phenotype appears to lie within the spectrum of a BMD-like disease, which likely represents an approximate ceiling of what would be expected of micro-dystrophin’s efficacy in the clinic.

**Figure 3.**
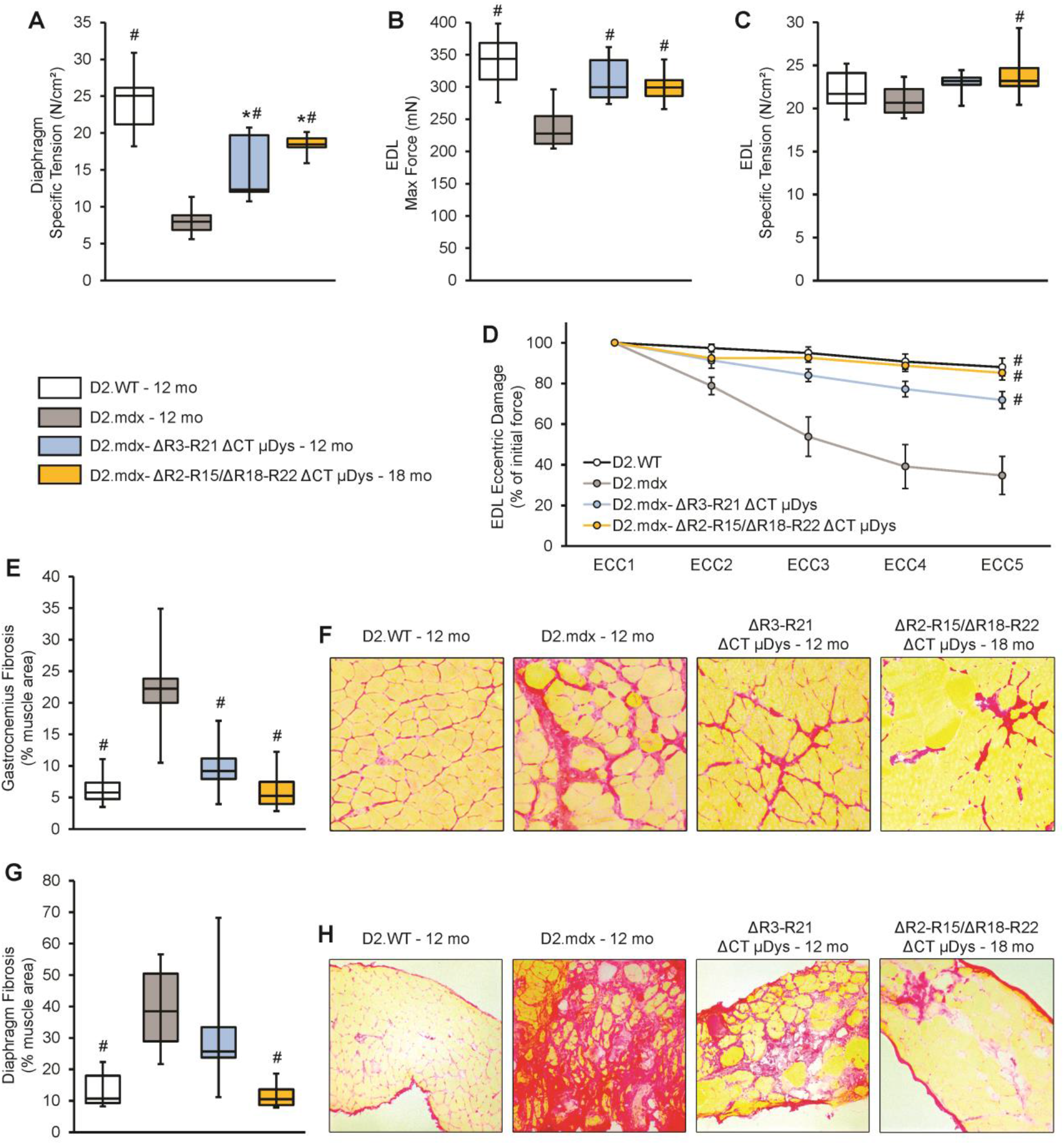
Micro-dystrophin provides partial rescue of D2.*mdx* skeletal muscle.\. Male D2.*mdx* mice were treated with micro-dystrophin (µDys) gene therapy at 1 month of age (mo; refer to **Figure 2a**). At the terminal endpoints of 12- and 18-mo, *ex vivo* muscle function was performed for the (**a**) diaphragm and (**b-d**) extensor digitorum longus muscles (EDL) of D2.WT, untreated D2.*mdx*, and µDys-treated D2.*mdx* (D2.*mdx*-µDys) mice (n = 7-10). (**f/h**) Representative picrosirius red-stained images of the gastrocnemius and diaphragm muscles and (**e/g**) accompanying fibrosis quantifications for these groups. *p < 0.05 vs. D2.WT values; #p < 0.05 vs. D2.*mdx* values.

### Micro-dystrophin gene therapy does not benefit D2.mdx hearts

We next turned our attention to the impact of micro-dystrophin gene therapy on the hearts of D2.*mdx* mice. Cardiac function was assessed by collecting electrocardiograms and echocardiograms at 6 and 12 months of age, with and additional assessment at 18 months for mice that received ΔR2-R15 ΔR18-R22 ΔCT micro-dystrophin.

As shown in the echocardiography data of **Figure 4A-G**, neither micro-dystrophin overtly impacted D2.*mdx* hearts at 6 months of age; however, untreated D2.*mdx* hearts also do not exhibit much difference in function from D2.WT hearts at this age. Electrocardiogram abnormalities arise prior to functional deficits in D2.mdx mice (as in DMD patients), and micro-dystrophin treatment did normalize T-wave duration at 6 months of age (**Figure 5C**), in agreement with a previous report (24).

**Figure 4.**
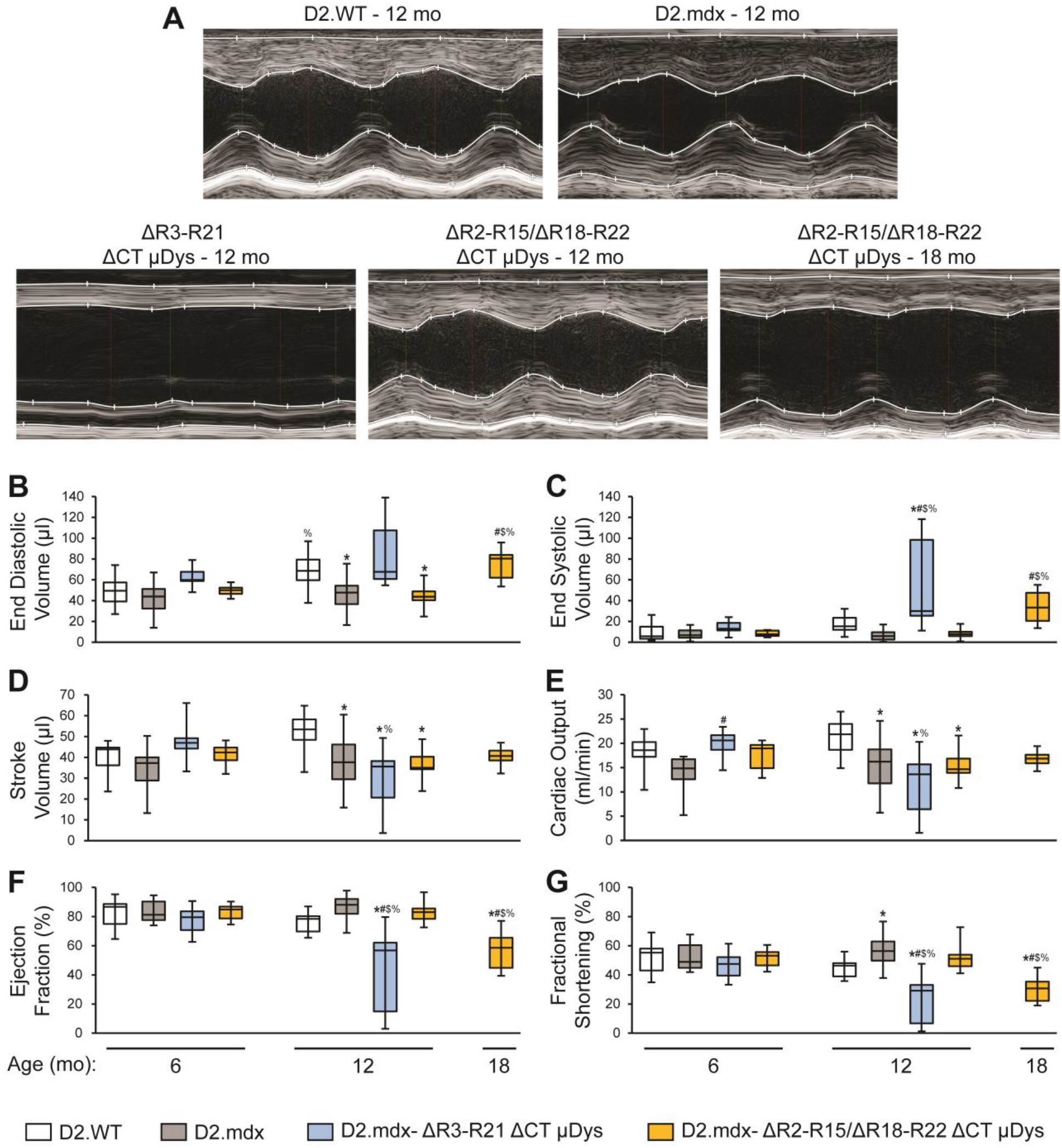
Long-term micro-dystrophin expression causes cardiomyopathy in D2.*mdx* mice. Male D2.*mdx* mice were treated with micro-dystrophin (µDys) gene therapy at 1 month of age (mo; refer to **Figure 2a**). (**a**) Representative left ventricle (LV) M-mode ultrasound images. Select (**b-g**) echocardiography parameters are shown, as measured at 6-12- and 18-mo (n = 7-23). *p < 0.05 vs. age-matched D2.WT values; #p < 0.05 vs. age-matched D2.*mdx* values; $p < 0.05 vs. 12-month-old D2.*mdx*-ΔR2-R15 ΔR18-R22 ΔCT values; %p < 0.05 vs. group-matched 6-month values.

**Figure 5.**
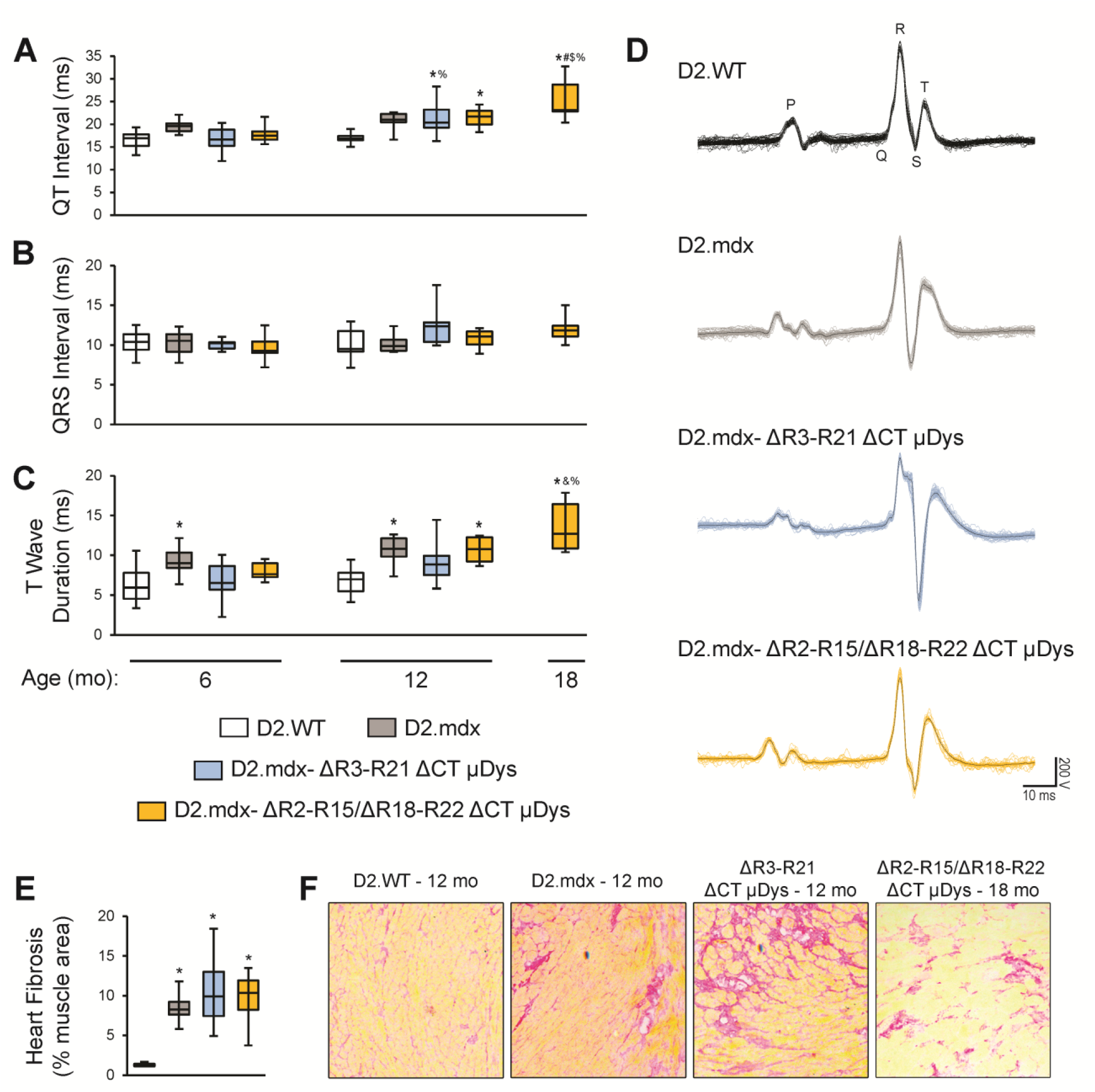
Electrocardiogram abnormalities and cardiac fibrosis persist with micro-dystrophin treatment. Male D2.*mdx* mice were treated with micro-dystrophin (µDys) gene therapy at 1 month of age (mo; refer to **Figure 2a**). Select (**a-c**) electrocardiogram parameters are shown, as measured at 6-12- and 18-mo (n = 7-14). (**d**) Representative electrocardiogram tracings (**f**) Representative picrosirius red-stained images of the heart and (**e**) accompanying fibrosis quantifications for these groups. *p < 0.05 vs. age-matched D2.WT values; #p < 0.05 vs. 12-month-old D2.*mdx* values. %p < 0.05 vs. group-matched 6-month values; $p < 0.05 vs. 12-month-old D2.*mdx*-ΔR2-R15 ΔR18-R22 ΔCT values; &p < 0.05 vs. 12-month-old D2.*mdx*-ΔR3-R21 ΔCT values.

A disturbing effect of the ΔR3-R21 ΔCT as compared to ΔR2-R15 ΔR18-R22 ΔCT micro-dystrophin treatment on D2.*mdx* hearts was found at 12 months of age. In contrast to untreated D2.*mdx* hearts and those treated with the ΔR2-R15 ΔR18-R22 ΔCT micro-dystrophin, which showed progression of diastolic dysfunction, as indicated by a decrease in LV EDV **(Figure 4B**), ΔR3-R21 ΔCT micro-dystrophin treated D2.*mdx* mice developed a severe cardiomyopathy. This micro-dystrophin-induced cardiomyopathy exhibits features of DCM, including marked reductions in LV systolic functional parameters of ejection fraction (EF; **Figure 4F**), fractional shortening (FS; **Figure 4G**), stroke volume (SV; **Figure 4D**), and cardiac output (CO; **Figure 4E**) as well as prominent enlargement of the LV chamber size, as indicated by an increase in EDV (**Figure 4B**). Pulsed-wave Doppler of blood flow through the mitral valve reveal that these mice also have reduced early and atrial blood flow ratios (E/A; **Supplemental Figure 3A**) and an increased isovolumic relaxation time (IVRT; **Supplemental Figure 3B**). These DCM-like features are clearly evident in the representative 12-month LV M-mode images found in **Figure 4A**. Additionally, 12-month-old ΔR3-R21 ΔCT micro-dystrophin treated D2.*mdx* mice displayed prolonged QT intervals as seen in **Figure 5A** and the ECG tracings in **Figure 5D**. D2.mdx mice that received ΔR2-R15 ΔR18-R22 ΔCT micro-dystrophin succumbed to a similar heart failure with reduced systolic function (EF and FS) and dilation of the LV (increased EDV) at 18 months of age. Unlike the findings in the gastrocnemius and diaphragm (**Figure 3E-H**), neither micro-dystrophin reduced the amount of cardiac fibrosis present in the D2.mdx mice (**Figure 5E/F**). These data indicate that AAV-micro-dystrophin treatment can potentially have a detrimental impact on the heart, depending on the design. However, note that the non-detrimental micro-dystrophin (ΔR2-R15 ΔR18-R22 ΔCT) had no discernable impact on the heart, and followed the same disease progression as untreated hearts.

### Potential mechanisms contributing to micro-dystrophin-induced cardiomyopathy

We sought to explore potential mechanisms contributing to these detrimental cardiac outcomes. These investigations have led us to suspect two potential causes of this micro-dystrophin-induced cardiomyopathy: **a)** micro-dystrophin competes with and displaces endogenously-expressed utrophin at the cardiomyocyte sarcolemma and **b)** the long-term overexpression of micro-dystrophin protein contributes to overload of the ubiquitin-proteosomal system (UPS), resulting in impairments in cardiomyocyte protein quality control. We present below the data and observations in support of the first mechanism (utrophin displacement) likely causing of micro-dystrophin-induced acceleration of cardiomyopathy with ΔR3-R21 ΔCT but not with ΔR2-R15 ΔR18-R22 ΔCT.

The heart normally expresses a combination of utrophin and dystrophin, with potentially overlapping and as well as specific roles that are yet to be clearly defined. The ability of these two orthologous proteins to link the cytoskeleton to the extracellular matrix through their interactions with common partners is consistent with some degree of functional redundancy. Indeed, utrophin protein levels in the heart increase in absence of dystrophin (26-28), and the removal of utrophin worsens the cardiac phenotype in the B10.*mdx* mice (29-31), with the total removal of utrophin being worse than haploinsufficiency. Thus, it is clear that utrophin can partially mitigate the loss of dystrophin. To potentially explain how high levels of micro-dystrophin leads to cardiomyopathy, we sought to determine if micro-dystrophin displaces utrophin from the cardiomyocyte membrane, as it is possible that strong overexpression of micro-dystrophin may phenocopy utrophin ablation via replacement with a truncated, and potentially less functional, dystrophin molecule. Therefore, we assessed the relative amounts of utrophin at the cardiac membrane by both immunofluorescence of tissue sections and immunoblotting of membrane enriched fractions of cardiac extracts from D2.WT, untreated D2.*mdx*, and micro-dystrophin-treated D2.*mdx* mice, in order to discern whether micro-dystrophin displaces utrophin to levels below those of D2.WT hearts. In both assays, the hearts of ΔR3-R21 ΔCT micro-dystrophin-treated mice exhibited significant decreases in utrophin immunoreactivity at the membrane on the order of ∼60% of D2.WT levels and ∼30% of D2.*mdx* levels (**Figure 6A-D**). In contrast, ΔR2-R15 ΔR18-R22 ΔCT micro-dystrophin treatment resulted in normalized utrophin levels. To test the clinical relevance of this finding, we compared the utrophin displacement of ΔR3-R21ΔCT micro-dystrophin to its hinge-3 containing clinical-counterpart, ΔR3-R19/ΔR20-R21ΔCT, and found that the two constructs displace utrophin to the same extent (**Supplemental Figure 4A/B**).

**Figure 6.**
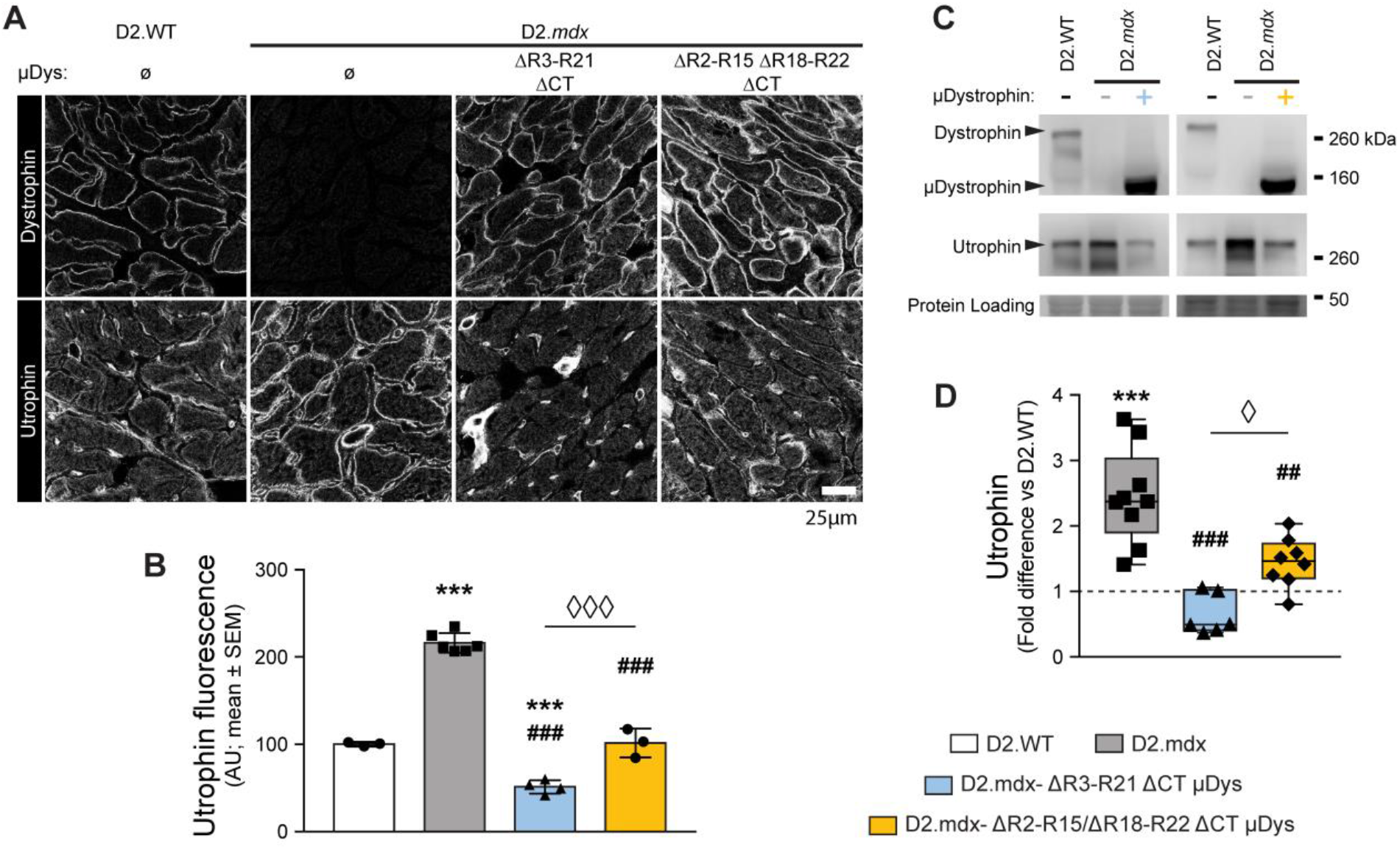
Micro-dystrophin overexpression displaces utrophin from cardiomyocyte sarcolemma. (A) Transverse sections of hearts from D2.WT, D2.*mdx* and micro-dystrophin-treated D2.*mdx* mice were labeled for dystrophin and utrophin. The increased sarcolemmal utrophin expression in absence of endogenous dystrophin is negated in cardiomyocytes over-expressing micro-dystrophins. (B) Utrophin immunouorescence intensity at cardiomyocyte membrane shows that utrophin sarcolemmal localization is increased approximately 2-fold in absence of dystrophin (d2.*mdx*), but reversed and even repressed (∼1/2 of D2.WT) upon AAV-mediated over-expression of µDys ΔR2-R15 ΔR18-R22 ΔCT (n=4) and µDys ΔR3-R21 ΔCT (n=3), respectively (One-way ANOVA; ### p<0.001 vs D2.WT, *** p<0.001 vs D2.mdx, ⧫⧫⧫ p<0.001 vs D2.mdx + μDys ΔR3-R21 ΔCT, Tukey post-hoc comparison). (C, D) Similarly, western blots of heart membrane-enriched samples reveal an approximately 2-fold upregulation of membrane-associated utrophin in D2.*mdx* (n=9) that is normalized or reduced to approximately 60% of the levels D2.WT hearts (grey dotted line) upon over-expression upon AAV-mediated over-expression of µDys ΔR2-R15 ΔR18-R22 ΔCT (n=6) and µDys ΔR3-R21 ΔCT (n=8), respectively (One-way ANOVA; ### p<0.001 vs D2.WT, *** p<0.001 vs D2.*mdx*, ⧫ p<0.05 vs D2.*mdx* + μDys ΔR3-R21 ΔCT, Tukey post-hoc comparison). Scale bars: 25 μm

This ability of ΔR3-R21ΔCT micro-dystrophin to outcompete utrophin for association with the sarcolemma is not restricted to cardiomyocytes, as its expression in D2.*mdx* skeletal muscle also results in utrophin displacement from the sarcolemma of muscle fibers (**Supplemental Figure 5A**). Sarcolemmal utrophin localization is maintained, however, in myofibers overexpressing ΔR3-R19/ΔR20-R21ΔCT microdystrophin, similar to what is observed in cardiomyocytes (**Figure 6**). Micro-dystrophin and utrophin, thus, display a complementary and mutually exclusive pattern of expression in both heart and skeletal muscles of micro-dystrophin-treated D2.*mdx* mice. This likely results from competition between the two proteins for common binding partners present within the sarcolemma. There are two sites of utrophin accumulation in wild type skeletal muscle fibers where utrophin, along with dystrophin, accumulates in high density: neuromuscular junction (NMJ) and myotendinous junction (MTJ). Utrophin accumulation at these specialized portions of myofibers appear unperturbed despite overexpression of either micro-dystrophin (**Supplemental Figure 5B/C**). The absence of any noticeable utrophin depletion by micro-dystrophins at NMJs (**Supplemental Figure 5A**) could result from the assembly of specialized sub-regions of the post-synaptic apparatus in which dystrophin (along with voltage-gated sodium channels) and utrophin (together with nicotinic acetylcholine receptors; nAChRs) are segregated (32-35) (34, 36, 37). Such organization suggests distinct interactions that recruit dystrophin and utrophin to their respective domains with specificity. The degree of micro-dystrophin overexpression in skeletal muscle achieved in these experiments (approximately 10-fold lower than in the heart) may be insufficient to overcome utrophin’s affinity to its interacting proteins at the NMJs. Alternatively, but not mutually exclusive, the sheer density of utrophin at NMJs and MTJs that appears to far exceed extrajunctional sarcolemma (Supplemental Figure 5B,C) could afford it continued association at the synapse. In case of NMJs, the density of nAChRs at the NMJs is measured to be up to a 1000-fold greater than at the extrasynaptic portions of the myofiber surface (38). while it is unknown whether utrophin levels at NMJs reach that of the nAChRs, its concentration at the synapse, as well as its potential to form protein interactions distinct from those of dystrophin, likely help maintain high-density synaptic accumulation of utrophin despite micro-dystrophin overexpression.

Another potential mechanism by which micro-dystrophin expression leads to cardiomyopathy, whether independently or in combination with utrophin displacement, is the saturation of the UPS by the excess micro-dystrophin molecules. Postmitotic cells, including cardiomyocytes, are especially susceptible to proteotoxicity stemming from accumulation of misfolded proteins, and impaired cardiomyocyte protein homeostasis has been shown to cause DCM-like cardiac phenotypes (39, 40). The sheer degree of overexpression (∼55 fold higher than endogenous dystrophin of wild-type hearts) may saturate the capacity of the cardiomyocytes to ensure that proteins maintain their functional conformation and to breakdown/recycle those that are misfolded or damaged. Accumulation of polyubiquitinated proteins can serve as a molecular signature for UPS saturation and can lead to cardiomyopathy by impairing both the proper clearing of damaged/misfolded proteins and the timely turnover of typically short-lived proteins with specific signaling or transcriptional roles (41, 42). Although we find a modest increase (∼20-40%) in polyubiquitinated (K48-linkage) proteins in the micro-dystrophin-treated hearts at the onset of dilated cardiomyopathy at 12-months in D2.*mdx* mice treated with the ΔR3-R21ΔCT micro-dystrophin and at 18-months with ΔR2-R15ΔR18-R22ΔCT, these values do not significantly differ from untreated D2.*mdx* hearts (**Supplemental Figure 6A/B**). However, when the degree of increased accumulation of polyubiquitinated proteins was plotted against cardiac ejection fraction from individual animals, a clear inverse correlation between the two emerged (Supplemental Figure 6C,D). This was true for both micro-dystrophins. At both, 12- and 18-months, micro-dystrophin treated hearts – tend to be with lowest ejection fractions – tend also to have highest accumulation of protein polyubuquitination. These data are consistent with the idea that disrupted protein homeostasis stemming from micro-dystrophin overexpression in cardiomyocytes may not be a primary cause of the dilated cardiomyopathy but may have accelerated the progression toward heart failure.

## DISCUSSION

Micro-dystrophin gene therapy clinical trials are currently underway for the treatment of DMD. In the current report, we sought to critically examine the long-term efficacy of two different micro-dystrophin gene therapies using a severe mouse model of DMD in order to better understand the impact and potential limitations of these emerging therapeutics for the treatment of DMD. Previously, we demonstrated that the DBA/2J background strain does not exhibit an inherent cardiomyopathy (43), validating it as a useful background strain for this study. Herein, we identified that long-term treatment of D2.*mdx* mice with AAV-packaged micro-dystrophins that are similar to two clinical versions results in widespread transduction of the skeletal muscle and slowing of, but not arresting, the progression of skeletal muscle disease. The treated muscles exhibit a slower progressing muscular degenerative disease, suggesting a conversion from DMD to a BMD-like pathology. Surprisingly, the clinical dose of one of the two different AAV-micro-dystrophins resulted in the development of a severe and early onset life-limiting dilated cardiomyopathic failure, while in the second case, there was extended life before experiencing a late onset DCM. The very different cardiac outcome despite similar impact in the skeletal muscle does not appear to be due to the function of the micro-dystrophin per se. The premature onset of this cardiomyopathy appears to be related to the degree of micro-dystrophin expression in the heart and the specific design of the micro-dystrophin that alters its competition with utrophin for binding to the dystrophin-glycoprotein complex. We provide evidence that micro-dystrophin expression at the levels achieved via high dose AAV delivery leads to displacement of native utrophin protein at the cardiomyocyte sarcolemma. The acceleration of the cardiomyopathy was due to a nearly complete displacement of utrophin, which only occurred for one of the two micro-dystrophins. These data highlight the benefits, limitations, and potential deleterious consequences, of maximizing micro-dystrophin overexpression in both skeletal and cardiac muscle for the treatment of DMD. Likely both micro-dystrophin constructs would show some benefit in the heart if the levels of expression did not result in utrophin displacement.

### Micro-dystrophin partially rescues skeletal muscle disease in D2.mdx mice

The potential to modify a severe DMD disease using a truncated dystrophin molecule was initially suggested by the existence of mildly-progressing BMD patients that express mutant dystrophin proteins missing most of the rod domain (44, 45). Therefore, the ultimate goal of micro-dystrophin gene therapy is to convert DMD into a milder disease. Indeed, we find that the long-term trajectory of the skeletal muscle phenotype of micro-dystrophin-treated D2.*mdx* mice does represent a milder dystrophy, with progressive pathology most notable in the diaphragm. This progressive myopathy does not appear to be due to loss of micro-dystrophin over time, as we initially anticipated, but rather is caused by the failure of micro-dystrophin to rescue all of the functions of full-length dystrophin, as in BMD.

It is likely that different designs of micro-dystrophin may slow the skeletal muscle disease to varying degrees. For instance, the micro-dystrophin ΔR2-R15/ΔR18-R22ΔCT construct is able to restore nNOS to the skeletal muscle membrane (24, 25), which may provide additional benefits to the muscle beyond sarcolemmal stability. Indeed, the diaphragm appears to be better rescued by this micro-dystrophin than by ΔR3-R21ΔCT (**Figure 3G/H**). This same region does not bind nNOS in the heart (46) but may serve other functions in the heart (47) and may have provided benefit that delayed the onset of DCM in the treated hearts, even though there was no impact on the onset of diastolic dysfunction (**Figure 4B**). On the other hand, the micro-dystrophin ΔR3-R21ΔCT construct may exhibit increased membrane binding by the inclusion of both repeats 1 and 2, which may enhance membrane localization and functional stability of the micro-dystrophin protein (48). Attempts to restore some or all of the missing C-terminus in order to better reconstitute the membrane complex may also improve function and further slow disease progression. However, it is becoming increasingly clear that all regions of dystrophin serve specific roles, thus, any micro-dystrophin is likely to be a physiological compromise in lieu of full-length dystrophin. Only animal models that recapitulate aspects of the human disease, such as the D2.*mdx* mouse, can reveal which compromises are likely the most efficacious for dystrophic muscle. Ultimately it is likely that other types of therapies will need to be combined with micro-dystrophin gene therapy for the optimal management of DMD.

### Micro-dystrophin overexpression causes cardiomyopathy

The most unexpected finding of this study is that long-term overexpression of the ΔR3-R21ΔCT micro-dystrophin in the heart leads to an early onset dilated heart failure, while the overexpression of the ΔR2-R15/ΔR18-R22ΔCT micro-dystrophin leads to a much later onset of dilated heart failure. While there are numerous pre-clinical publications evaluating the efficacy of systemic AAV-mediated delivery of micro-dystrophin, many of these studies did not assess cardiac function (25, 49-51). Of the studies that did evaluate cardiac function (via EKG and pressure-volume catheters), a dose less than is used in the current study (and current clinical trials) was used and/or the short treatment length would have prevented them from observing a progression to heart failure (24, 52-55). Therefore, to our knowledge, this is the first study that has assessed the long-term cardiac function of a severe mouse model of DMD following micro-dystrophin gene therapy using the high dose of AAV being used in clinical trials.

We propose that no micro-dystrophin is necessarily detrimental, and should in fact be beneficial, to the heart unless it is expressed at very high levels and/or it displaces utrophin. It is possible that the overexpression of the micro-dystrophin contributes to the accelerated onset of heart failure, based in the correlation between accumulation of polyubiquitinated proteins and ejection fraction (**Supplemental Figure 6)**. High level of transgene overexpression, *per se*, can be detrimental if the increased protein turnover overloads the protein breakdown capacity of the cell (39) and likely puts an extra energetic load on an already stressed heart (56). Indeed, it was previously shown that 100-fold transgenic overexpression of a mini-dystrophin was associated with cardiac toxicity (57). This mini-dystrophin is likely more efficacious in the heart than any micro-dystrophin, which may allow higher levels of overexpression to be tolerated.

The dose of AAV (2 × 10^14^ gc/kg) used for this study was chosen to mirror doses being used in all of the ongoing clinical trials with AAV-micro-dystrophin in DMD patients (23). The choosing of this dose for the clinical trials has been dictated by the attempt to transduce as many skeletal muscle fibers as possible, which is assessed by post-injection muscle biopsies (23). There has been no consideration, however, of what this dose escalation may mean for the heart, and adequate modeling of these high doses and their long-term impact on the heart has not been previously performed. Furthermore, these efforts have incorporated the use of promoters that drive high level expression of the transgene in both skeletal and cardiac muscles, with the assumption that all micro-dystrophins will benefit both muscle types regardless of how much overexpression is achieved. The degree of cardiac expression achieved in preclinical models and DMD patients is dependent on the efficiency of cardiac muscle infection of the AAV capsid serotype used and the strength of the promoter in the heart. Based on the differential amounts of transgene expression we (**Figure 2**) and others (51) have noted between murine skeletal and cardiac muscles, it is reasonable to assume that the heart is receiving more vector per cell than the skeletal muscle fibers. Since all of the promoters being used in the clinical trials have been optimized for expression in both muscle types, a similar phenomenon may be occurring in the clinic. A recent update on one clinical trial reported promising gene therapy transduction and micro-dystrophin expression in the skeletal muscles of trial participants 1 year following treatment (23). It is not clear if AAV transduction or promoter activity differs in the human hearts, compared to those achieved in mouse hearts, and, thus, the level of cardiac micro-dystrophin expression that is being achieved in DMD patients is unknown. If the preclinical expression differential translates into humans, these patients will also exhibit high expression levels in the heart. As older patients, who have fewer intact skeletal muscles, more muscle fibrosis, and more body fat than younger patients, are treated, the danger is that even greater amounts of virus will be directed to the heart, leading to still greater levels of micro-dystrophin overexpression

Furthermore, the displacement of utrophin by a potentially less functional micro-dystrophin may further accelerate heart failure. It is clear that cardiac disease progression is greatly slowed by the high degree of utrophin expression in the heart (29, 30), therefore, displacement of utrophin by a less functional micro-dystrophin expressed at high levels may exacerbate the cardiomyopathy. In this study, equivalent cardiac overexpression was achieved, but only one of the two micro-dystrophins tested displaced utrophin from the cardiomyocyte membrane, which was accompanied by an accelerated disease progression to dilated heart failure. From our data, it does not appear that all micro-dystrophin constructs will compete with utrophin equally well, or if some will provide greater functionality and thus be a more successful replacement for utrophin. How well a specific micro-dystrophin substitutes for utrophin or full-length dystrophin will depend on which regions are in the micro-dystrophin and which regions are most critical for proper cardiac function. Competition will likely depend not only on the degree of overexpression, but also on the design of the micro-dystrophin. The third micro-dystrophin in clinical trials, shown in Figure 1 (ΔR4-R23ΔCT), has also shown to express at much higher levels in the heart than skeletal muscles of mice (51). What impact this will have will likely depend on the extent that it competes with utrophin. Of note, this construct contains repeats 1-3 which have been shown to have higher lipid-binding affinity than repeats 1-2 (48) and therefore may better outcompete utrophin.

A recent study (58) demonstrated that micro-dystrophin is beneficial in the total absence of utrophin in a B10.*mdx* background, slowing disease progression to a degree that was comparable to utrophin haploinsufficiency. However, comparing that study with this study is difficult since the utrophin was missing throughout development, possibly allowing adaptations that cannot occur with acute postnatal displacement of utrophin by overexpression of micro-dystrophin. It is also not clear how much overexpression of micro-dystrophin was achieved in the study of Howard *et al*. (58). Given the much slower disease progression in the C57 background, it is possible that lack of rescue of the utrophin-deficient C57Bl10.mdx hearts may not be evident until a more advanced age than was studied (12 months of age). In the absence of utrophin, we would predict that all micro-dystrophin constructs should slow the onset of cardiac dysfunction and failure as compared to no intervention.

### Conclusion

It appears that the acceleration of cardiac disease progression we observed following high dose systemic delivery of AAV-ΔR3-R21ΔCT-micro-dystrophin is primarily due to loss of utrophin and replacement by a truncated dystrophin that is less functional. High-level protein overexpression in cardiomyocytes may further accelerate the progression to failure. On the other hand, delivery of the same dose of AAV-ΔR2-ΔR15/ΔR18-R22ΔCT-micro-dystrophin resulted in similar levels of micro-dystrophin expression but did not reduce utrophin levels in the cardiomyocytes below wild type levels. In this case, the rescue of the skeletal muscle prolonged life until there was a late onset DCM. To determine how much benefit the heart derived from the ΔR2-ΔR15/ΔR18-R22ΔCT micro-dystrophin treatment, we would need to compare this treatment group to mice with skeletal muscle rescue without any micro-dystrophin expression in the heart, which we currently cannot do.

Whether or not the DMD patients currently being dosed with high doses (2-3 × 10^14^ gc/kg) of AAV-micro-dystrophin in clinical trials are at risk of accelerated cardiac disease, is also unclear. It may be years before we can address this question, given that it requires 8-12 months to clearly see this cardiomyopathy development in mice. However, our data suggest that there is reason to be concerned that while the skeletal muscles may be improved in the DMD patients receiving the current AAV-micro-dystrophin vectors, the hearts may not be improved, and possibly even be worsened by these treatments. Even if the treatment is somewhat beneficial, the increased load on the heart due to the improved skeletal muscle function may accelerate the onset of DCM and failure. Therefore, frequent monitoring of the cardiac status of these patients should be performed and prophylactic use of cardio-protective drugs should be considered. If the observations in this study are recapitulated in DMD patients, then delivery of micro-dystrophin to the heart may need to be dissociated from skeletal muscle via the use of promoters designed to be much weaker in the heart than in skeletal muscle.

## METHODS & MATERIALS

### Animals

This study used male D2.WT (Jax# 000671) and D2.*mdx* (Jax# 013141) mice from colonies originally obtained from Jackson Laboratory. Mice were housed 1-5 mice per cage, randomly assigned into groups, provided *ad libitum* access to food (NIH-31 Open formulation diet; Envigo #7917), water, and enrichment, and maintained on a 12-hour light/dark system. All animal studies were approved and conducted in accordance with the University of Florida IACUC.

### Micro-dystrophin constructs and vector production

Codon-optimized µDys was synthesized by Genscript (Piscataway, NJ) and cloned into a pAAV shuttle plasmid containing the striated muscle-specific CK8 promoter (20) and a minimized synthetic polyadenylation signal sequence (59). AAV viral vector packaging was performed using the triple-transfection method, as previously described (21, 60).

### Ex vivo muscle function

Maximal tetanic tension assessments of the EDL and diaphragm muscles were evaluated as previously described (61) by the University of Florida Physiological Assessment Core. Subsequently, a series of 5 eccentric contractions with (stimulated at 80 Hz for 700 ms) a stretch of 10% optimal length was imposed on the muscle in the last 200 ms of each contraction. Each contraction was separated by a 5-minute rest period. Following experimental procedures, muscles were weighed, frozen embedded in OCT or snap-frozen, and stored at -80 C until further use.

### Echocardiography and electrocardiograms

Electrocardiograms and transthoracic echocardiograms were performed using the Vevo 3100 pre-clinical imaging system (Fujifilm Visualsonics). Mice were anesthetized using 3% isoflurane and maintained at 1.5-2% to keep heart and respiration rates consistent among treatment groups. Body temperature was maintained at 37°C throughout imaging. Electrocardiograms were imported into LabCharts (ADInstruments) for analysis. Four images were acquired for each animal: B-mode parasternal long axis (LAX), B-mode short axis (SAX), M-mode SAX, and apical four-chamber view with color doppler and pulsed-wave doppler. M-mode SAX images were acquired at the level of the papillary muscle. Flow through the mitral valve was sampled at the point of highest velocity, as indicated by aliasing, with the pulsed-wave angle matching the direction of flow. Images were imported into Vevo LAB for analysis. Measurements of M-mode SAX and pulsed-wave doppler images were made from three consecutive cardiac cycles between respirations.

### Fractionation, protein extraction, and immunoblotting

Snap-frozen mouse heart and gastrocnemius muscles were finely crushed and homogenized in a phosphate-based homogenization solution – 2 mM sodium phosphate, 80 mM NaCl, 1 mM EDTA (62) – supplemented with 1 mM PMSF, phosphatase/protease inhibitor cocktail (ThermoFisher Scientific), and centrifuged at 12,000 x g for 10 min at 4°C. The supernatant (soluble cytosolic fraction) is collected. The pelleted non-cytosolic (including membrane and cytoskeletal) fraction is then re-suspended in the extraction buffer [homogenization solution supplemented with the following: 20 µg/ml DNase I (Sigma), 10 µM Vinblastine (Caymen Chemicals), 100 mM Swinholide A (Caymen Chemicals), 100 mM Mycalolide B (Focus Biomolecules), 1% Digitonin (Biosynthe), 0.5% NP-40, 1% SDS] and extracted on-ice for 45-min with occasional vortexing, followed by a 15-min incubation at 37°C. The insoluble fraction was pelleted by centrifugation at 12,000 x g for 10 min at 4°C, and soluble membrane fraction was collected. The protein concentration of soluble cytoplasmic and membrane fractions was determined using the Bio-Rad Protein Assay (Bio-Rad Laboratories). Samples were boiled in 4X sample buffer, proteins separated using a 4-12% SDS polyacrylamide gels (ThermoFisher Scientific) and transferred to nitrocellulose membranes using the iBlot system (Life Technologies). Membranes were incubated at room temperature with 5% BSA-TBST, then overnight with primary antibodies at 4°C. Following TBST washes and species-appropriate horseradish peroxidase-conjugated secondary antibody (Cell Signaling), incubated with ECL reagent (ThermoFisher Scientific), and imaged using the Li-Cor C-DiGit imaging system (Li-Cor Biosciences). Membranes were probed for GAPDH for cytosol/non-cytosol fractionation and stained with Ponceau S to control for equal protein loading and for normalization. The following primary antibodies were used for immunoblotting in the present study: MANHINGE1B (1:100; Clone 10F9; Developmental Studies Hybridoma Bank (DSHB)), MANEX1011B (1:100; Clone 1C7; DSHB), MANEX1011C (1:100; Clone 4F9; DSHB), utrophin-A (1:1000; ABN1739; EMD Millipore), Dystrobrevin (1:1000; #610766; BD Biosciences); Syntrophins (1:2000; #11425; Abcam), β-dystroglycan (1:500; #11017-1-AP; Protein Tech), Polyubiquitin (K48-linkage; 1:2000; #4389, Cell Signaling), and GAPDH (1:2000; SC-25778 ; Santa Cruz). Band signal intensities were measured using Image Studio Lite software (Li-Cor Biosciences), normalized to sample loading (Ponceau S stain), and reported relative to respective control samples.

### Immunofluorescence and histological evaluations

Fresh-frozen OCT-embedded hearts and gastrocnemius muscles were section at 10 µm and fixed in ice-cold acetone. The sections were re-hydrated in PBS, blocked in 5% BSA-PBS at room temperature and incubated with primary antibodies overnight at 4°C. Mouse tissue sections to be incubated with mouse monoclonal antibodies were first incubated with a solution containing donkey anti-mouse IgG AffiniPure Fab fragments (1:25 in PBS; Jackson ImmunoResearch #715-007-003) for one hour prior to blocking. Following PBS washes, sections were incubated at room temperature with species- and isotype-appropriate fluorescent dye-conjugated secondary antibodies and coverslipped using Prolong Gold anti-fade mounting medium (ThermoFisher Scientific). The following primaries were used for immunofluorescence in the present study: MANHINGE1B (1:100; Clone 10F9; DSHB), MANEX1011B (1:100; Clone 1C7; DSHB), MANDAG2 (1:100; Clone 7A11; DSHB), utrophin-A (1:1000; ABN1739; EMD Millipore), utrophin (1:50; VP-U579; Vector Laboratories), Dystrobrevin (1:500; #610766; BD Biosciences); Syntrophins (1:2000; #11425; Abcam), and α-sarcoglycan (1:50; VP-A105; Vector Laboratories). NMJs were identified using fluorescent dye-conjugated α-bungarotoxin (1:500; ThermoFisher Scientific) to label nAchRs localized to the postsynaptic motor endplates. Image acquisition was performed with a Leica Application Suite X software on either a Leica TSC-8 confocal system or a Leica DMR epifluorescence microscope equipped with a Leica DCF480 digital camera. Comparative images were stained, imaged, and processed simultaneously under identical conditions.

Picrosirius Red (PSR) staining was performed as previously described (13) following decalcification of muscle sections using Formical-2000 (StatLab). Slides were visualized with a Leica DMR microscope, and images were acquired using a Leica DFC310FX camera interfaced with Leica LAS X software. Images were processed and analyzed by investigators blinded to study groups using ImageJ software.

### Statistical analysis

Statistical analysis was performed using unpaired, two-tailed Welch’s T-test (α = 0.05), ANOVA (one-way, two-way, or repeated measures) followed by Tukey HSD post-hoc tests (α = 0.05), and Kaplan-Meier estimator analyses (α = 0.05), where appropriate. A P value less than 0.05 was considered significant. Data are displayed as mean ± SEM, box-and-whisker plots, or survival curves.

## ACKNOWLEDGMENTS

This work was funded by a Wellstone Muscular Dystrophy Cooperative Center grant (P50-AR-052646) from the NIH to HLS and DWH, a Parent Project Muscular Dystrophy grant to HLS. GG is supported by grants from NIH (R01NS076991-01, 1P01AI100263-01, 4P01HL131471-02, UG3 HL147367-01, and R01HL097088). Michael Matheny, Heejae Chun, and Lillian Wright are thanked for their technical support related to this project.

## AUTHOR CONTRIBUTIONS

Study design was contributed by CCH, YL, DWH, and HLS. Experimental procedures and data acquisition were conducted by CCH, YL, and DWH. Essential reagents were produced and provided by JX and GG. All authors were involved in data analysis, interpretation, data presentation, and manuscript writing.

## Conflict of Interested Statement

The authors have declared no conflict of interest exists.

## Data Availability

The datasets generated during and/or analyzed during the current study are available from the corresponding author on reasonable request.

## SUPPLEMENTAL FIGURES

**Supplemental Figure 1.**
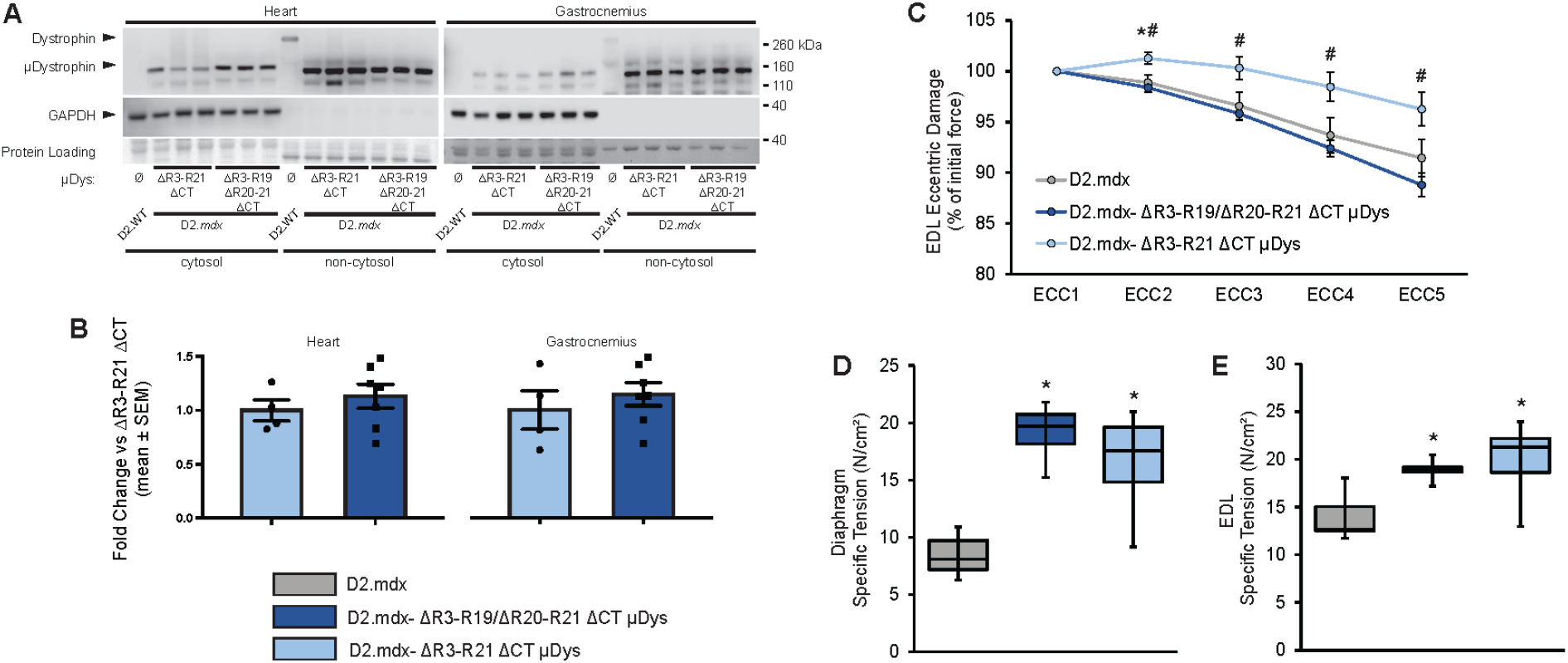
Equivalent expression and function of micro-dystrophins ΔR3-R21ΔCT and ΔR3-R19/ΔR20-R21ΔCT. (A,B) Western blots of heart and gastrocnemius samples were performed to compare the expression levels of the two micro-dystrophins – ΔR3-R21ΔCT and ΔR3-R19/ΔR20-R21ΔCT. For both micro-dystrophins (n=3 each), majority of the protein product was found in the membrane-enriched (non-cytosol) fraction as is the case with endogenous dystrophin protein (D2.WT). Total protein levels of the two micro-dystrophins did not differ in either tissue examined. Functionally, micro-dystrophin ΔR3-R21ΔCT provide treated muscle better protection against eccentric contraction-induced damage (C), while specific tension measured from both diaphragm (D) and EDL (E) both improved equally with either micro-dystrophins. *p < 0.05 vs. age-matched D2.mdx values; #p < 0.05 vs. age-matched D2.mdx-ΔR3-R19/Δ20-R21 ΔCT values.

**Supplemental Figure 2.**
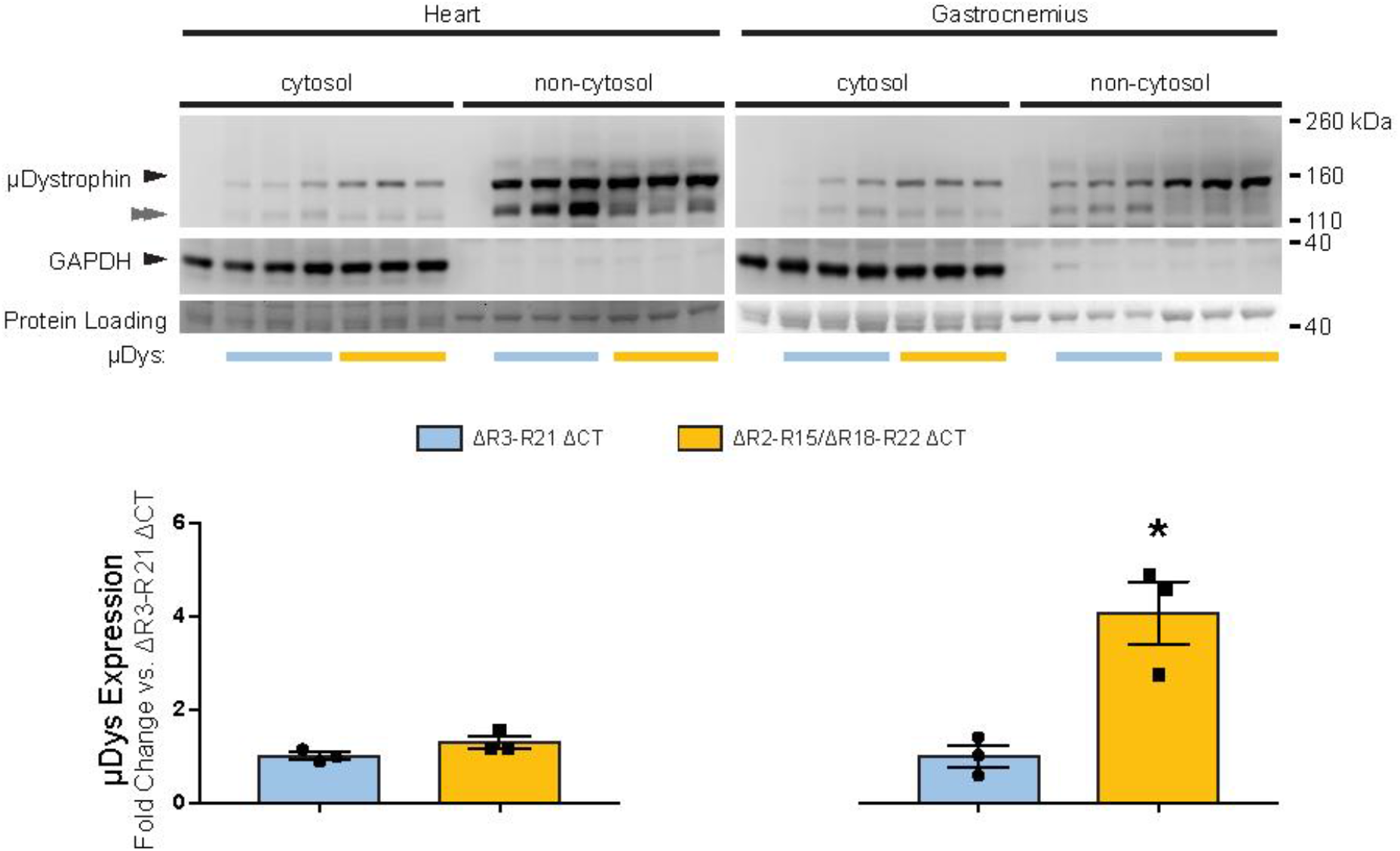
AAV-mediated striated muscle expression of micro-dystrophins ΔR3-R21ΔCT and ΔR2-R15 ΔR18-R22 ΔCT. Western blot analysis of heart and gastrocnemius samples from D2.*mdx* mice treated with either micro-dystrophins show robust expression of micro-dystrophins that preferentially target to membrane-enriched (non-cytosol) fraction. *p < 0.05 vs. D2.mdx-ΔR3-R19/Δ20-R21 ΔCT values.

**Supplemental Figure 3.**
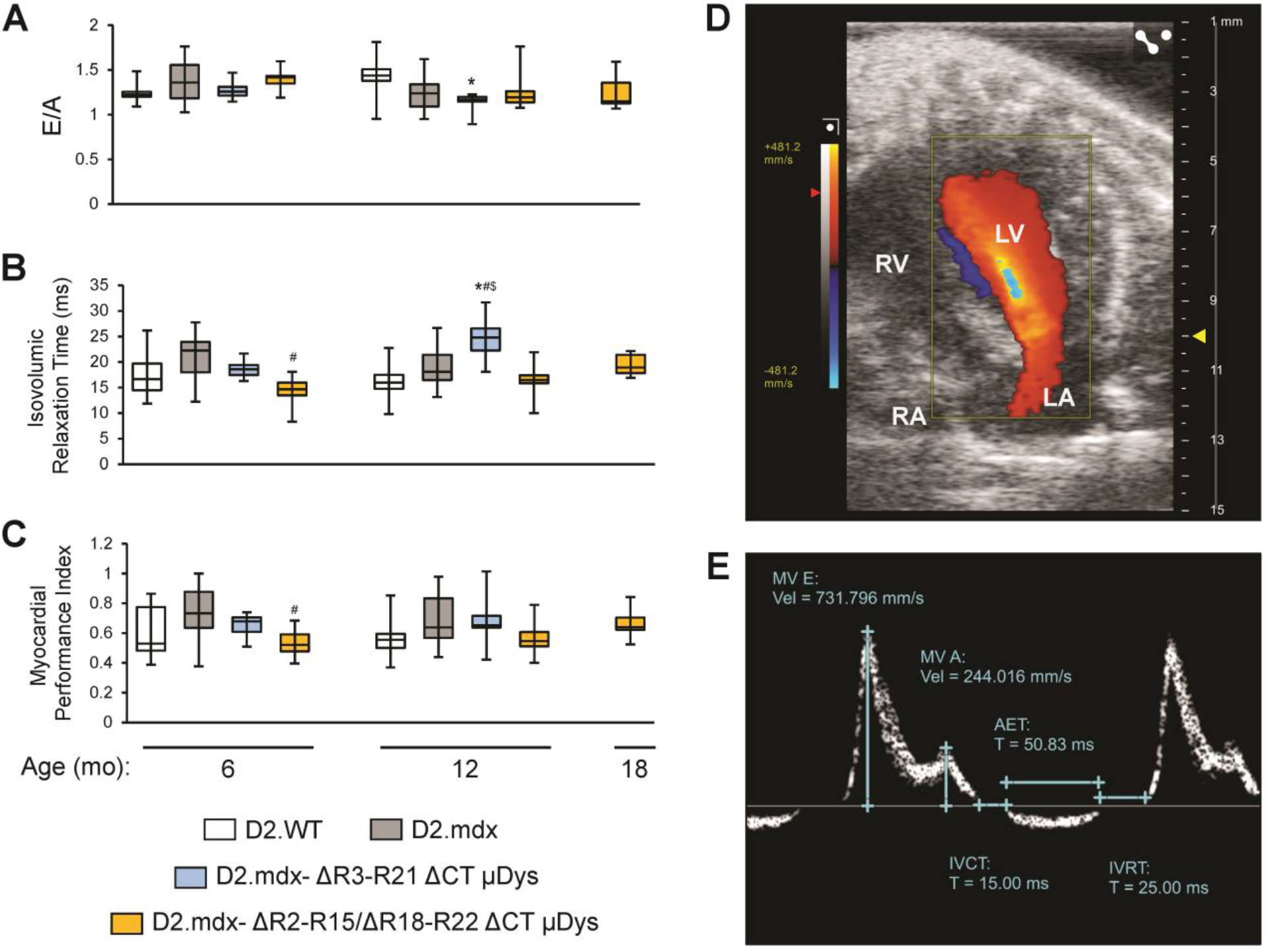
Diastolic dysfunction persists with micro-dystrophin treatment. Male D2.*mdx* mice were treated with micro-dystrophin (µDys) gene therapy at 1 month of age (mo; refer to **Figure 2a**). Select (**a-c**) pulsed-wave Doppler parameters are shown, as measured at 6-12- and 18-mo (n = 6-22). (**d**) Representative color Doppler image taken in the four chamber apical view and (**f**) representative pulsed-wave Doppler; *p < 0.05 vs. age-matched D2.WT values; #p < 0.05 vs. age-matched D2.*mdx* values; $p < 0.05 vs. 12-month-old D2.*mdx*-ΔR2-R15 ΔR18-R22 ΔCT values

**Supplemental figure 4.**
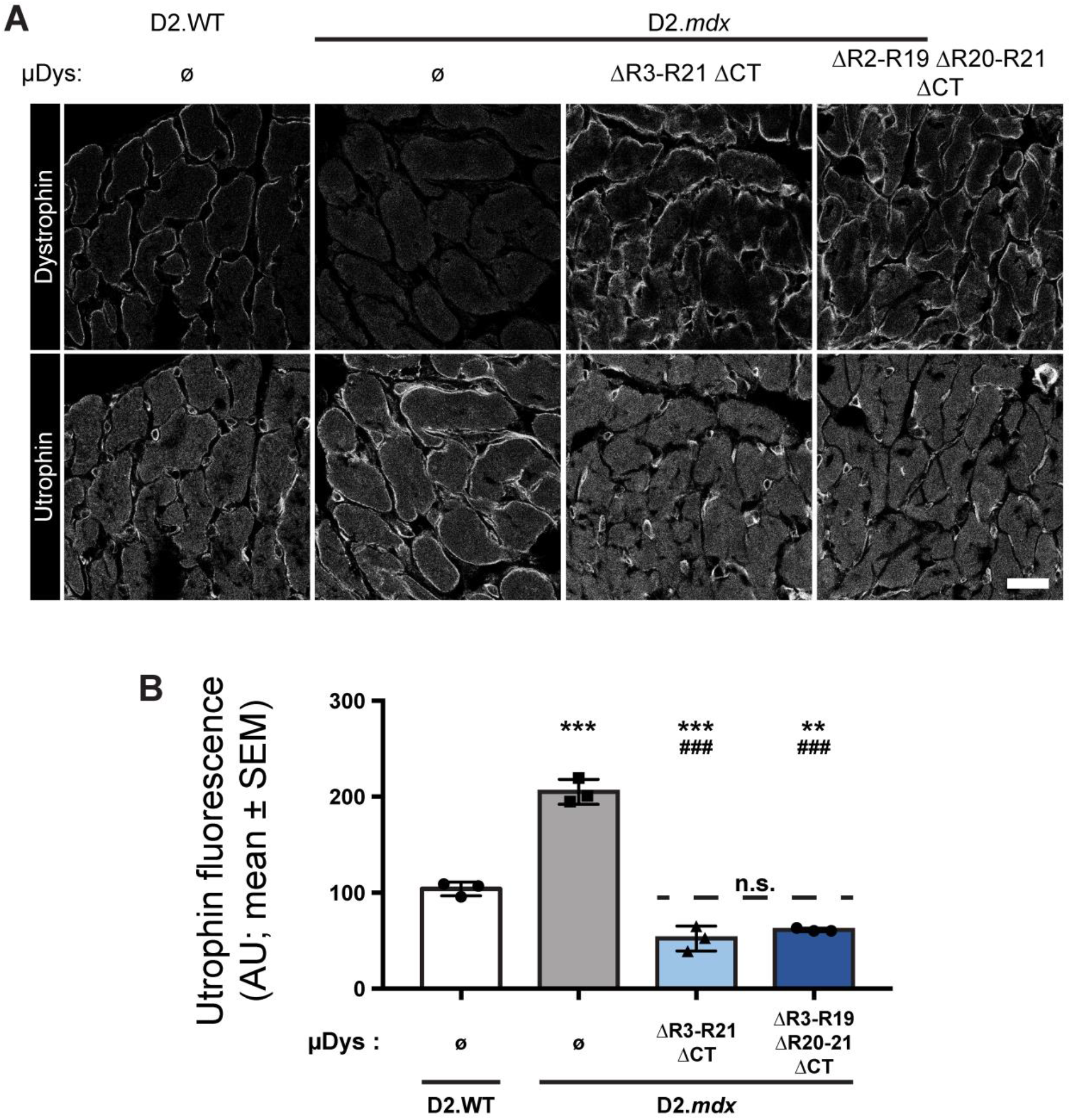
Equivalent utrophin displacement from sarcolemma in cardiomyocyte that express micro-dystrophins ΔR3-R21ΔCT and ΔR3-R19/ΔR20-R21ΔCT. (A) Transverse sections of hearts from D2.WT, D2.*mdx* and micro-dystrophin-treated D2.*mdx* mice were labeled for dystrophin and utrophin. (B) Both micro-dystrophins – ΔR3-R21ΔCT and ΔR3-R19/ΔR20-R21ΔCT – equally repressed sarcolemmal utrophin localization below levels found in D2.WT cardiomyocytes (n=3 each, One-way ANOVA; ### p<0.001 vs D2.WT, *** & ** p<0.001 and p<0.01 vs D2.mdx, Tukey post-hoc comparison). Scale bar: 25 µm.

**Supplemental figure 5.**
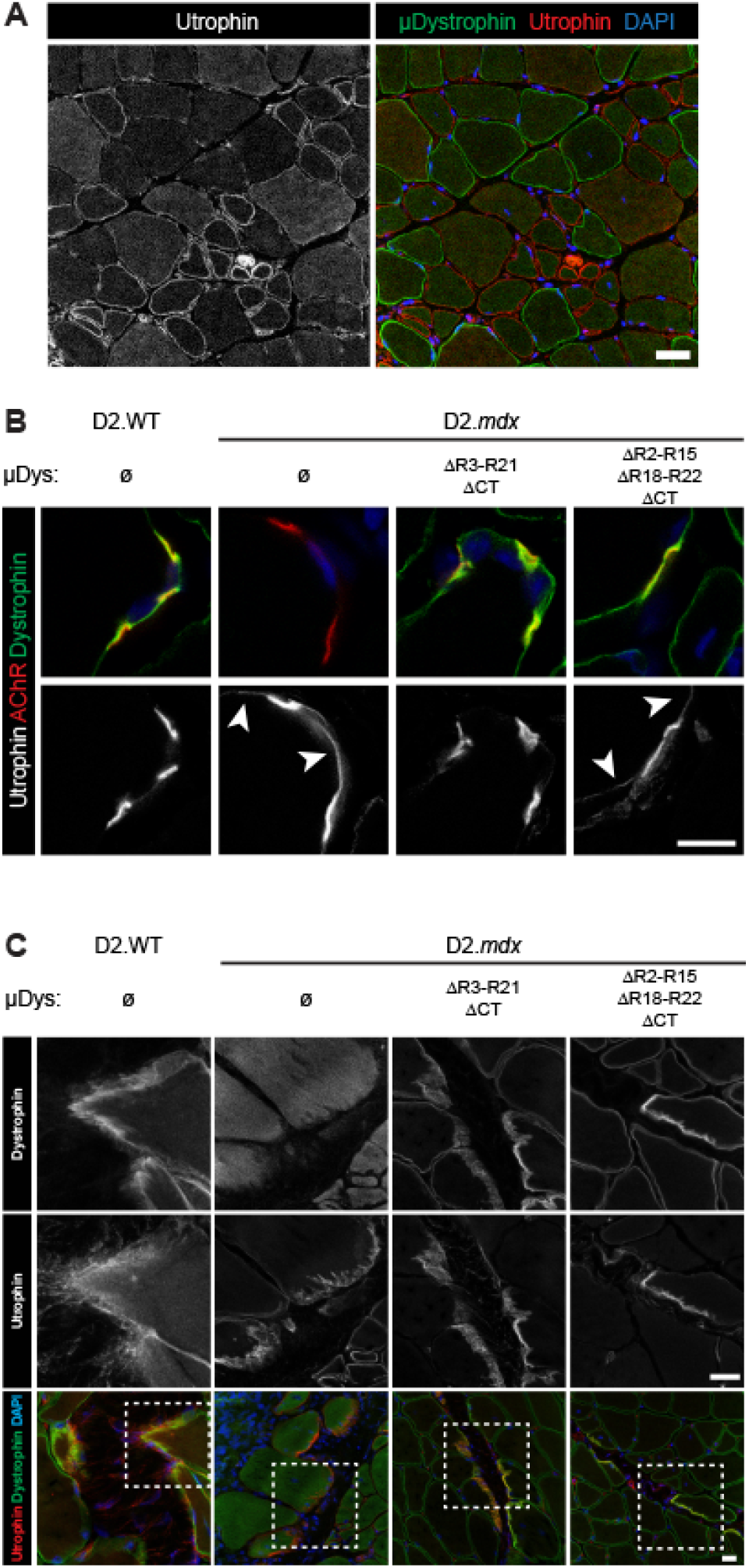
Micro-dystrophin overexpression in skeletal muscie fibers displaces utrophin from sarcolemma outside of neuromuscular and myotendinous junctions. (A) Micro-dystrophin and utrophin demonstrate a mutually exclusive pattern of sarcolemmal labeling amongst skeletal muscle fibers in mice treated with micro-dystrophin ΔR3-R21ΔCT. (B) Utrophin distribution is restricted to neuromuscular junctions in myofibers of D2.WT mice, but extends beyond the postsynaptic membrane (white arrowhead) in D2.*mdx* skeletal muscle fibers. The synaptic localization of utrophin (as indicated by co-localization with postsynaptic acetylcholine receptors (AChR) appears unperturbed by expression of either micro-dystrophins. Micro-dystrophin ΔR3-R21ΔCT, unlike ΔR3-R19/ΔR20-R21ΔCT, displaces extrasynaptic sarcolemmal utrophin (white arrowhead) in treated skeletal muscle fibers. (C) Utrophin remains concentrated at myotendinous junctions of muscle fibers overexpressing either micro-dystrophin. Scale bars: 10 µm (a, b); 20 µm (c)

**Supplemental figure 6.**
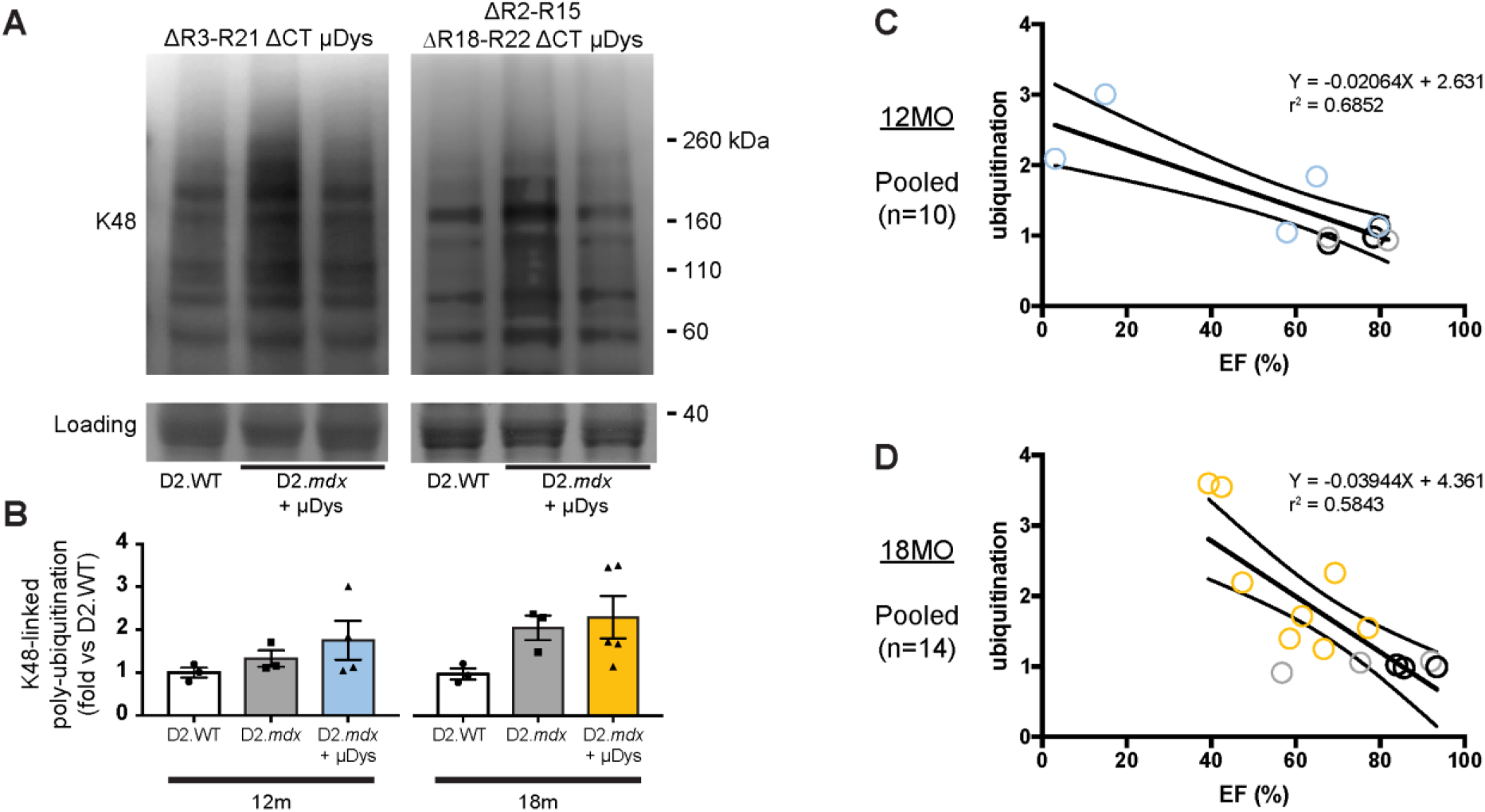
Cardiac ejection fraction inversely correlates with accumulation of polyubiquitinated proteins in cardiac lysates. (A) AAV-mediated over-expression of either micro-dystrophin protein does not lead to significant increase in the accumulation of polyubuiquitinated proteins in cardiac lysates. (B) A linear regression analysis demonstrates that an inverse relationship exists between cardiac ejection fraction and abundance of polyubiquitinated proteins (relative to age-matched D2.WT). With each age group, the slope of the best-fit line deviates significantly from zero (p<0.01).

## Notes

### Competing Interest Statement

The authors have declared no competing interest.

